# Induction of pluripotent oncogenic stem cells from mouse fibroblasts

**DOI:** 10.1101/2024.11.06.622204

**Authors:** Snehal Shabrish, Naorem Leimarembi Devi, Shubham Mohanty, Rohit Kumar, Relestina Lopes, Nabila Akhter, Naveen Kumar Khare, Gorantla V Raghuram, Vishalkumar Jadhav, Sushma Shinde, Pratik Chandrani, Indraneel Mittra

**Author notes:** Authors for correspondence: Indraneel Mittra Pratik Chandrani.

## Abstract

Natural biological agents that can transform normal somatic cells into cancer stem cells, have not been identified. We earlier reported that cell free chromatin particles (cfChPs) that circulate in blood of cancer patients can horizontally transfer themselves to healthy cells to induce dsDNA breaks and inflammation. Here we show that a single cell clone D5 developed from NIH3T3 mouse fibroblast cells treated with cfChPs isolated from sera of cancer patients exhibited upregulation of stemness related transcription factors and surface markers, and the ability to form spheroids in appropriate culture medium. Transcriptome analysis revealed upregulation of cancer related pathways including invasion, metastasis and stemness. Subcutaneous inoculation into SCID mice resulted in development of malignant tumors which expressed all three germline markers. Our results suggest the cfChPs that circulate in blood of cancer patients are oncogenic and can transform susceptible somatic cells into cancer stem cells with the potential to promote metastasis.

**Graphical Abstract:** 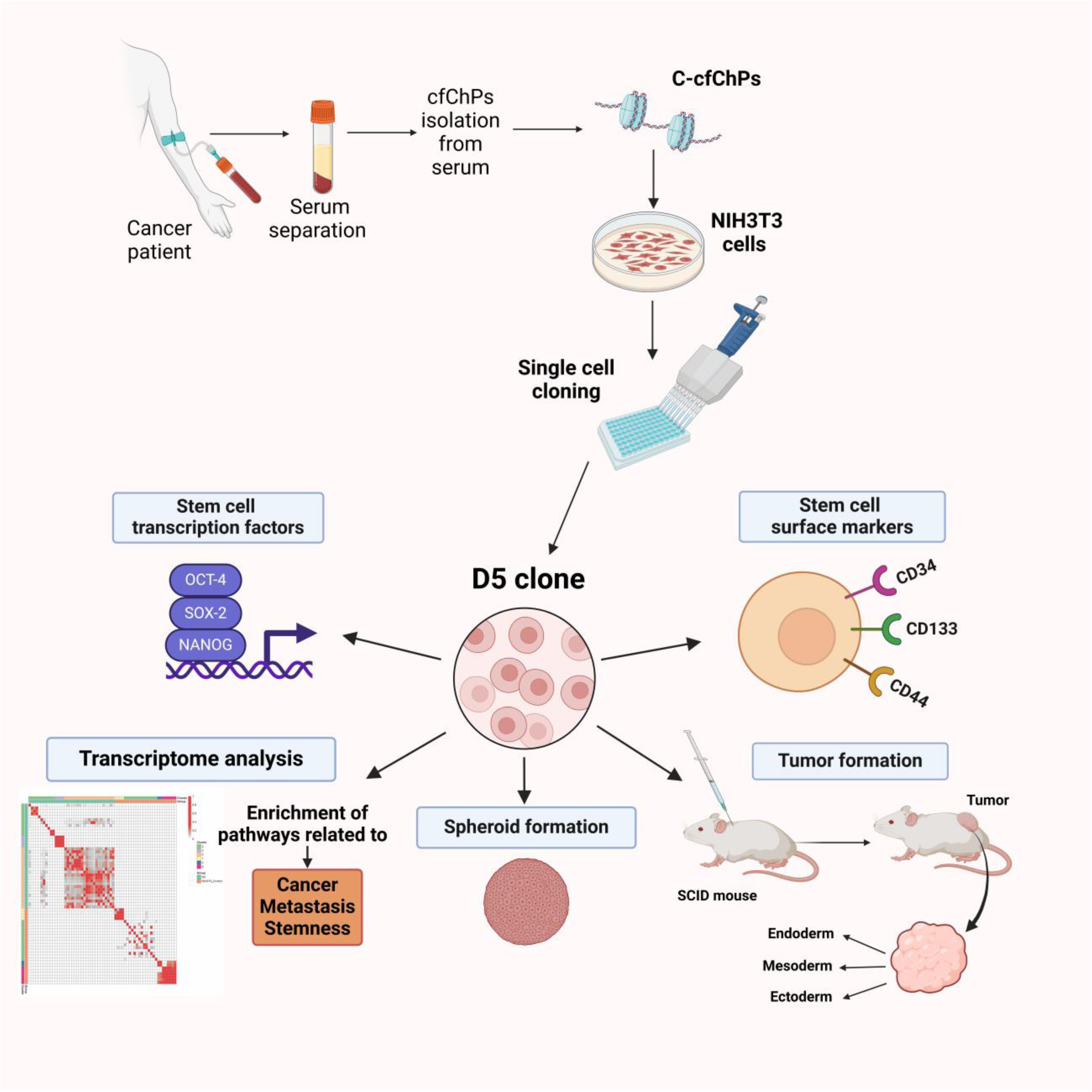

## Introduction

How cancer begins and metastasizes to distant organs are questions central to cancer research. Several theories of carcinogenesis have been proposed, of which the somatic mutation theory is the most dominant^1–3^. Cancer is thought to be the culmination of progressive stepwise accumulation of mutations in cancer causing genes^4,5^. Mutations in critical “driver” genes are thought to confer selective survival and growth advantage to cells leading to sustained cellular proliferation, invasion and metastasis^3,6^. Aberrant expression - both increased constitutive expression of oncogenes and repression or loss of tumour suppressor genes - forms the basis of the somatic gene mutation theory^7^. However, recent research has uncovered presence of tumour specific driver mutations in cells of normal tissues of healthy individuals^8,9^, and in case of the esophagus, more driver mutations were detected in normal epithelia than in the corresponding tumours^10^. Analysis of sequencing data from over 33 tumour types and over 9000 tumour samples detected driver mutations in only a handful of genes, suggesting the gene mutation may not be the primary cause of carcinogenesis in majority of cases^11^. Similarly, a recent report suggests that loss of the critical tumour suppressor gene p53 alone may not be sufficient for tumour progression^12^.

Another significant concept in cancer biology is stemness, which refers to a cell’s intrinsic capacity to exhibit characteristics associated with embryonal stem cells, including differentiation potential and self-renewal^13^. Cancer stem cells are a subpopulation of cells within a tumour which have the capacity for self-renewal, sustained proliferation, promoting carcinogenesis, inducing metastatic spread and causing tumor heterogeneity^13,14^. Tumors use stemness as fuel for their proliferation, invasion and resistance to treatment^13,14^. There have been a number of theories to explain the genesis and maintenance of pluripotent cells within tumors. They have included the cancer stem cell theory^15^, dedifferentiation theory^16^, somatic cell reprogramming theory^17^ and microenvironmental influence theory^18^. However, none of the above theories have been experimentally demonstrated to bring about stemness in tumours, and the physiological trigger for pluripotency remains to be discovered^19^.

In summary, in spite of extensive research, it has not been possible to identify a specific trigger for the carcinogenic process or for the induction of cancer stemness. It has been suggested that there are yet undiscovered mechanisms governing these processes which have been lost in the vast and complex reductionist science of cancer research^20^.

We have earlier reported that cell-free chromatin particles (cfChPs) that circulate blood of cancer patients, or those that are released locally from dying cancer cells, can readily enter into healthy cells to illegitimately integrate into their genomes leading to dsDNA breaks and activation of apoptotic and inflammatory pathways^21,22^. As DNA damage and inflammation are two critical hallmarks of cancer^23–25^, their concurrent activation may be a potential stimulus for oncogenic transformation of the affected cells. In the present study we investigated whether a single cell clone developed over a decade earlier from NIH3T3 mouse fibroblast cells that had been treated with cfChPs isolated from pooled sera of cancer patients would show cancer related molecular phenotypes, especially those related to pluripotent cancer stem cells.

## Results

### Morphological changes in D5 cells overtime

We observed that the cellular morphology of the D5 clone which was developed in 2011 changed over time. While in initial stages the cells were rounded, refractile and loosely attached to the culture dish, they gradually changed their morphology to become spindle shaped and adhered to the plastic surface, albeit still appearing distinctly different from the NIH3T3 cells from which they were derived (Supplementary Figure S1a). When we performed FISH analysis on metaphase spreads prepared 13 years after the D5 clone was developed, i.e. in 2024, human DNA representing cfChPs were still detectible and were found to be associated with the chromosomes. This finding suggested that cfChPs had continued to be involved in the oncogenic process since the inception of the D5 cells in 2011 (Supplementary Figure S1b).

### Expression of stem cell related transcription factors by D5 cells

qRT-PCR analysis of D5 cells revealed upregulated stem cell related transcription factors NANOG, SOX-2 and OCT-4. The degree of activation of all three markers ranges between 2-3 fold (Figure 1a). Analysis of D5 cells by flow cytometry detected expression of the stem cell related surface markers CD34 and CD133. However, the expression of CD44 was not statistically different from NIH3T3 control cells (Figure 1b). The flow cytometry gating strategy is given in Supplementary Figure S2.

**Figure 1.**
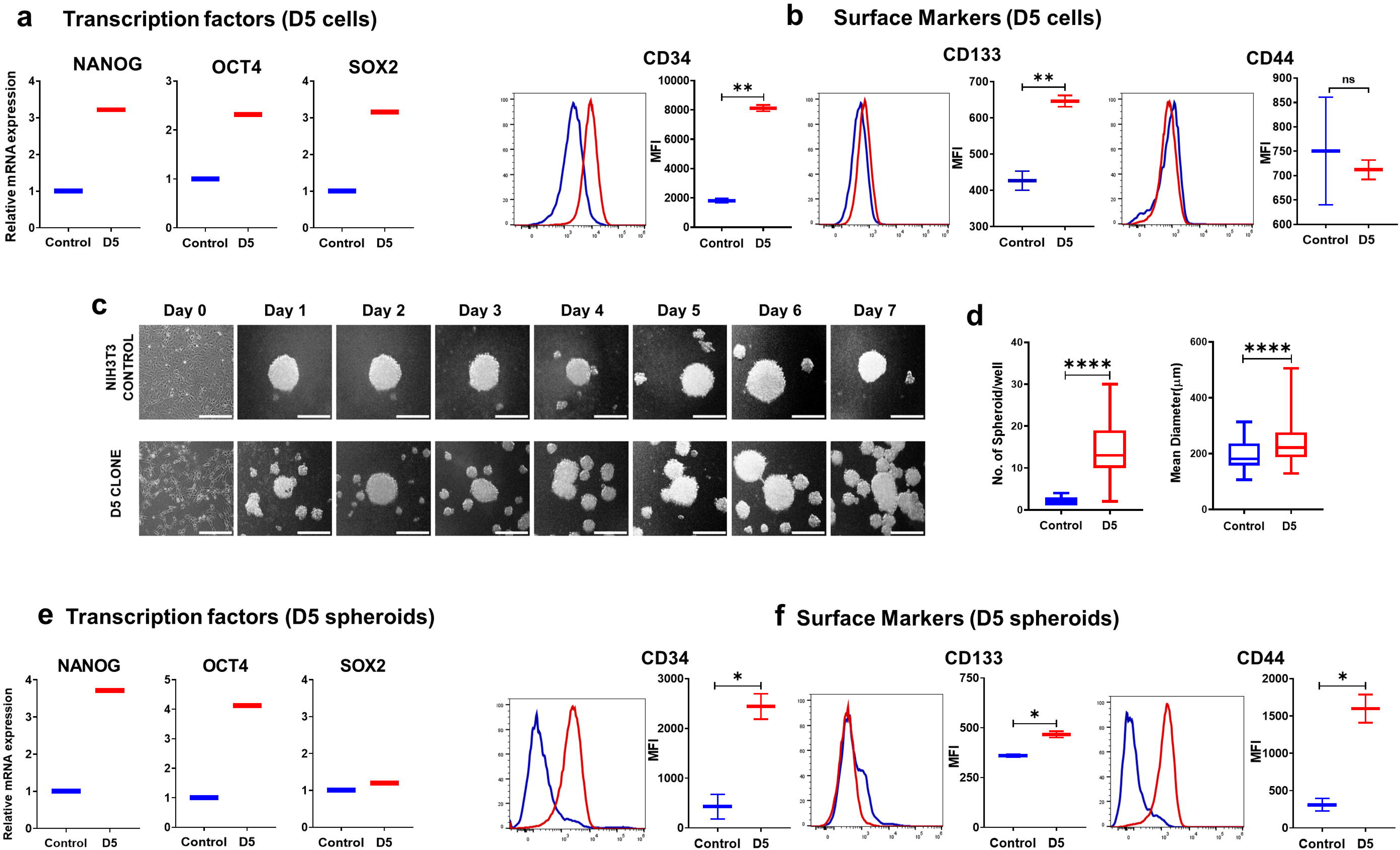

### Spheroid formation by D5 cells

When D5 cells were grown on ULA plates in stem cell medium, spheroid formation was visible from day 2 onwards with their growth accelerating in a time-dependent manner (Figure 1c). A quantitative analysis on day seven revealed highly significant differences in both number and size of the spheroids generated by D5 cells compared to control NIH3T3 cells (p< 0.0001) (Figure 1d).

### Expression of stem cell related transcription factors by D5 spheroids

Spheroids of NIH3T3 and D5 cells were trypsinized and subjected to qRT-PCR and flow cytometry analysis. The former revealed significantly increased expression of the transcription factors, viz. NANOG and OCT-4 but not of SOX-2 (Figure 1e). Flow cytometry detected significantly increased expression of all three stem cell related surface markers viz. CD34, CD133 and CD44 (Figure 1f). The flow cytometry gating strategy is given in Supplementary Figure S2.

### Detection of cancer and stem cell related pathways by transcriptome analysis

#### RNA-seq quality validation

Two replicates each for D5 and NIH3T3 samples were sequenced for whole transcriptome with a median coverage of 30 million reads (Range: 24 – 34 million) (Supplementary Table S1). FastQC was utilised to assess mean per-base quality scores and per-sequence quality scores of RNA-seq data. This analysis indicated a Phred quality score above 30 for all samples, affirming the high quality of RNA-seq reads (Figure 2a & Supplementary figure S3a). Multidimensional scaling (MDS) plot revealed a clear separation between NIH3T3 control and D5 samples (Figure 2b). MDS plot showed that D5 samples clustered together, indicating high consistency across all genes, however, there was marked scattering for control samples. Cook’s distance distribution was visualized as a boxplot that demonstrated comparable distribution across all samples (Supplementary figure S3b).

**Figure 2.**
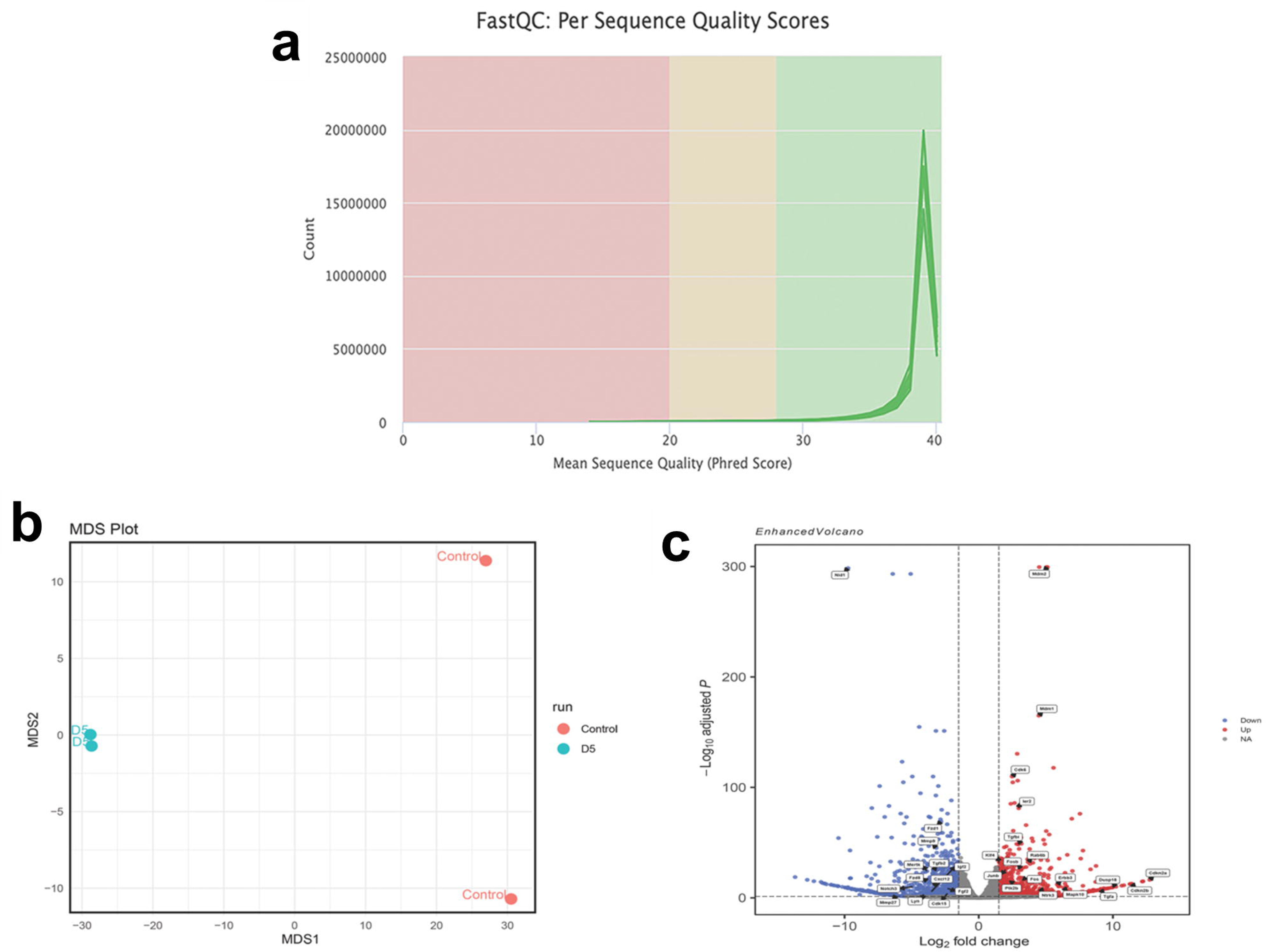

#### Identification of differentially expressed genes (DEGs)

DESeq2 analysis identified a total of 1573 DEGs, of these, 772 genes were upregulated, exhibiting a Log2 Fold Change (Log2FC) > 1.5 and an adjusted p-value (padj) < 0.05. The genes that were downregulated numbered 801 exhibiting a Log2FC <-1.5 and a padj <0.05. The heatmap representing the expression pattern of DEGs is shown in Supplementary Figure S3c, illustrating consistent expression patterns across samples. A complete list of DEGs in D5 compared to control samples is tabulated in Supplementary Table S2. Cancer associated genes such as Kruppel-like transcription factor 4 (Klf4), cyclin-dependent kinase inhibitor 2A (Cdkn2a), cyclin-dependent kinase inhibitor 2B (Cdkn2b), transforming growth factor alpha (Tgfa), mitogen-activated protein kinase 10 (Mapk10), cyclin-dependent kinase 6 (Cdk6), and transformed mouse 3T3 cell double minute 2 (Mdm2) were observed to be upregulated (Figure 2c). Conversely, the most significant downregulated genes include frizzled class receptor 1 (Fzd1), matrix metallopeptidase 9 (Mmp9), transforming growth factor, beta 2 (Tgfb2), frizzled class receptor 8 (Fzd8), C-X-C motif chemokine ligand 12 (Cxcl12), notch 3, fibroblast growth factor 2 (Fgf2), and insulin-like growth factor 2 (Igf2) as illustrated in (Figure 2c).

#### Identifying Significantly Enriched Pathways Through GSEA

GSEA analysis detected alterations in the expression of biological pathways associated with D5 samples. Pathway databases such as Hallmarks, Wikipathways, Biocarta and Reactome for enrichment analysis resulted a list of 36 upregulated pathways with normalized enrichment score (NES)>1.5 and a nom pval< 0.05 (Supplementary table S3). We observed a positive enrichment of pathways that are associated with cancer and cancer stem cell characteristics such as CTCF pathway, ERBB signaling pathway, P53 pathway, Apoptosis pathway, ATM pathway, Cellular senescence pathway and Oncogene induced senescence pathway (Figure 3a & Supplementary figure S4). Additionally, we observed 26 downregulated pathways in D5 samples, indicated by NES <-1.5 and a nom pval < 0.05 (Supplementary table S4). The selected GSEA plot for downregulated pathways is shown in (Supplementary figure S5).

**Figure 3.**
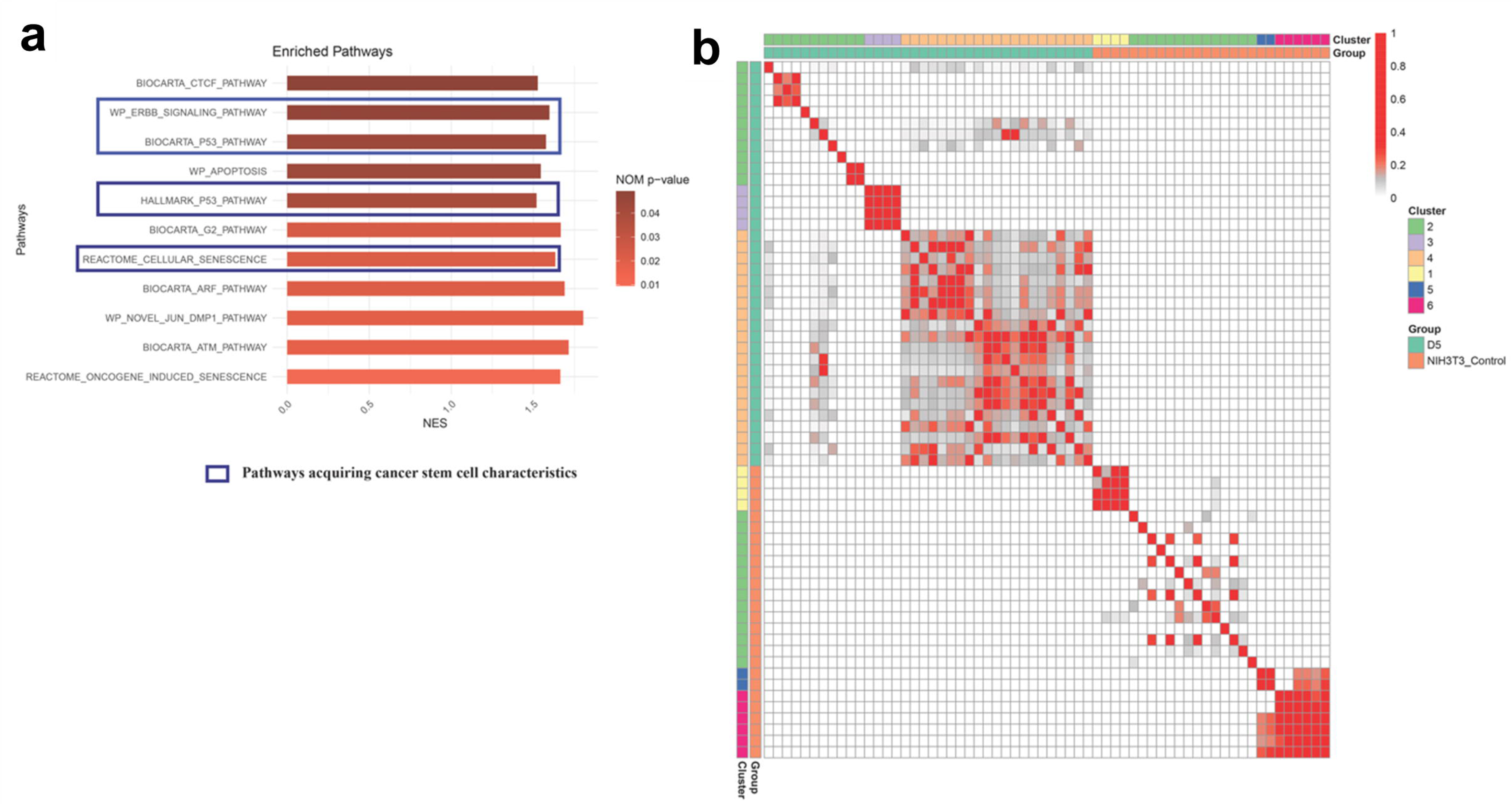

Three clusters for positive enriched pathways were identified based on shared genes (Figure 3b). The largest clusters included CTCF, Jun DMP1, Trafficking of AMPA receptors, P53, ATM, G2, ARF, Apoptosis, regulation of TP53, G1 to S DNA damage and senescence pathways. Additionally, key genes observed in this cluster include Mdm2, Cdkn1a, Cdkn2a, Rb1, Mapk1 and Ubc. The second cluster comprised of metabolism-related pathways and featuring significant genes such as Aldoa, Pgam1, Eno1 and Pgk1. The third cluster involved pathways that are associated with integrin and GPER1 signaling. Two clusters were observed for negatively enriched pathways. The first cluster was related to cell-cell communication, cell junction organization, and cell-extracellular matrix interactions while the second encompasses electron transport pathways (Supplementary figure S6).

### D5 cells form tumors in SCID mice

The NOD SCID mouse inoculation study was conducted twice: in 2018 and in 2024. In both studies 1 X 10^6^ NIH3T3 control cells and clone D5 cells were inoculated subcutaneously. In both experiments tumours developed in the D5 inoculated animals (5/5 in the first experiment and 8/8 in the second experiment). The rate of growth of the tumours was robust and palpable tumours developed within 2-3 weeks following inoculation (Figure 4a). No tumours developed in the NIH3T3 control inoculated mice (0/5 in the first experiment and 0/9 in the second experiment) after observing for tumour development for 10 weeks. Hematoxylin and eosin (H&E) stained FFPE sections of the tumours were indicative of fibrosarcoma (Figure 4b). FISH analysis of the formalin fixed paraffin embedded (FFPE) sections of the tumours using a human specific whole genomic probe revealed the presence of human DNA in the tumours indicating that cfChPs derived human DNA had persisted since the development of D5 clones 13 years earlier (Figure 4c).

**Figure 4.**
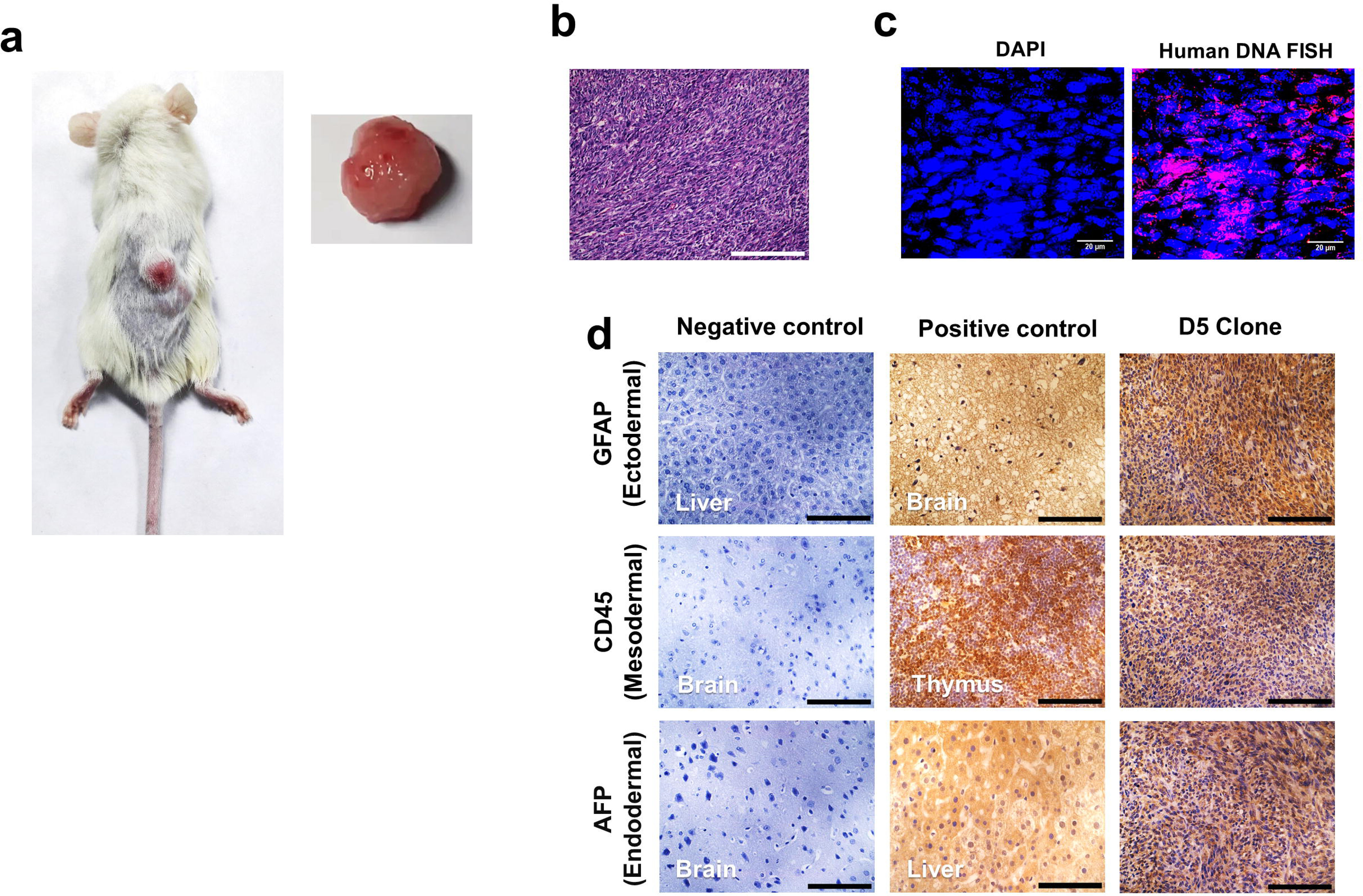

### D5 induced tumors express germline markers

We next performed immuno-histochemistry (IHC) analysis on the FFPE tumour sections to characterize the tumour cells and identify their origin. We detected expression of all three germline markers viz. ectoderm (GFAP), mesoderm (CD45) and endoderm (AFP) indicating that the D5 derived tumours were pluripotent (Figure 4d).

## Discussion

We have demonstrated in this article that cfChPs that circulate in sera of cancer patients have oncogenic potential and can transform NIH3T3 mouse fibroblast cells into cancer stem cells. The fact that clone D5 had retained its oncogenic and pluripotent properties for 13 years since it was developed, during which time the cells had undergone multiple freeze-thaw cycles, suggests that acquisition of these properties was not transitory and that the D5 clone is a stable population of cancer stem cells. This suggestion is supported by our demonstration that the D5 cells could rapidly form pluripotent tumours when inoculated into SCID mice 13 years after the cells were developed.

We employed NIH3T3 cells to develop the D5 clone as these cells are pre-disposed to undergo transformation in response to oncogenic stimuli^26^. NIH3T3 cells have been extensively used in transformation assays which had led to the discovery of oncogenes^27,28^. Significantly, the classical oncogene experiments had used microgram quantities of DNA, and CaPO4 precipitation or other transfection agents to introduce foreign DNA into host cells^29,30^. Incorporation of DNA resulted in development of transformed foci with low efficiency which appeared after several weeks of culture^31^. On the other hand, the D5 clone was developed following treatment of NIH3T3 cells with as little as 10ng of cfChPs, and since cfChPs can be readily internalized by healthy cells^21^, no transfection agents were required. These findings lead us to propose that cfChPs are natural oncogenic agents that circulate in blood of cancer patients and are the essential triggers for the development of pluripotent cancer stem cells.

The mechanism(s) by which cfChPs derived from sera of cancer patients transformed the NIH3T3 cells into cancer stem cells need to be addressed. In this context, a recent report by us based on an experimental design identical to that used to develop the D5 clone may be informative^32^. NIH3T3 cells were treated with cfChPs (10ng) isolated from sera of cancer patients and were serially passaged. We discovered that, once internalized by NIH3T3 cells, the cancer derived cfChPs displayed unusual intracellular processes and functions. The multiple internalized cfChPs, which comprised of DNA sequences that were wildly different from one another, had randomly joined together to create complex concatemers, some of which appeared to be larger than several megabase pairs.

Numerous tasks carried out by the concatemers were similar to the nuclear genome. They could replicate themselves within the NIH3T3 cells and orchestrate the synthesis of RNA, RNA polymerase, ribosomal RNA, ribosomal proteins and many other human proteins that appeared as intricate multi-peptide fusion proteins. Significantly, the concatemers harbored human LINE-1 and Alu transposable elements, which amplified and expanded in copy number over time in culture. The concatemers synthesized the enzymes reverse transcriptase and transposase which allowed them to rearrange themselves within the mouse genome causing repeated damage to the host cell DNA. Based on these findings we hypothesized that a cell harbors two autonomously functioning genomes: one that is inherited (hereditary) genome and numerous others that are acquired (predatory) genomes. Given that the “predatory” genomes can produce a wide variety of new proteins, including multiple oncogenes, and act as transposable element, suggested that their presence have oncogenic and evolutionary implications^32^. The extensive and diverse genetic changes that were detected in our transcriptome analysis of the D5 cells (discussed below) with enriched pathways relating to both oncogenesis and pluripotency may be the consequences of the anarchical activities of the “predatory” genomes within the NIH3T3 cells.

Transcriptome analysis detected upregulation of the stemness-related gene Klf4, known as one of the Yamanaka factors, and has been reported to be critical to the induction of pluripotent stem cells^33^. We observed several pathways with cancer stem cell-like characteristics such as ERBB signaling pathway, cellular senescence (reverse sequence) pathway and p53 pathway to be upregulated in D5 cells. In studies by others, these pathways were found in cancer cells that acquire stem-cell-like properties^34–36^. The upregulation of the ERBB pathway is significant as the ERBB signaling is known to regulate the stemness and differentiation status of cells^37^. The ERBB pathway influences downstream pathways viz. PI3K/AKT, which has been shown to maintain stemness and inhibit differentiation in embryonic stem cells^34,38^. While senescence is often considered to be a tumor-suppressing mechanism by preventing damaged or pre-cancerous cells from dividing, it can paradoxically promote stem-like properties in surviving cancer cells^36,39,40^. We observed an upregulation in the senescence pathway highlighting the potential for its stemness-promoting effects. Additionally, Lee et al.^35^ reported a notable link between the p53 pathway and Wnt signaling in mouse embryonic stem cells (mESCs). Their research, employed ChIP-on-chip and gene expression microarrays, which revealed that p53 directly activates the Wnt pathway, significantly inducing the expression of several Wnt ligands, including Wnt8b, in response to both genotoxic and non-genotoxic stress. This finding corresponds with our results showing enriched p53 signaling and upregulation of Wnt8b, suggesting a unique mechanism in mESCs that helps maintain stem cell properties under stress. The authors propose that this p53-mediated activation of Wnt signaling plays a critical role in preventing differentiation thereby maintaining the stem cell population.

Our study also identified several oncogenic pathways such as ATM pathway, CTCF pathway, and apoptosis pathway. Although, the ATM pathway typically acts as a tumor suppressor, its upregulation in cancer cells has been linked to oncogenic effects such as enhanced survival, proliferation, and metastasis^41^. Furthermore, a multifunctional transcription factor CTCF has been reported to play a role in the onset of various cancers, including breast cancer^42^, hepatocellular cancer^43^, and colorectal cancer^44^.

The findings of our study have implications for cancer metastasis by suggesting that cfChPs derived from dying cancer cells that circulate in blood can transform susceptible host cells to form new cancers in distant organs that could masquerade as metastasis. Such a model circumvents many of the problematic steps of the existing model that have not been fully elucidated^45,46^. For example, the new model bypasses the contentious issue of epithelial mesenchymal transition to explain how cancer cells exit the tumour parenchyma to enter the circulation^47^? The new model is also not answerable to the puzzling questions as to how extravasated cells withstand the shearing force of circulation^48^, evade immune attack^49^, navigate capillaries with diameters of ∼5 µm^50^, when the tumour cells themselves measure >20 µm^45,50^ and cross the blood-brain-barrier^51^.

Our study may be criticized on the ground that our findings based on NIH3T3 cells may not be relevant to the real world of cancer metastasis in humans. NIH3T3 cells are karyotypically unstable^26^ and are uniquely pre-disposed to readily undergo transformation in response to oncogenic stimuli, and that such unstable cells may not exist in the human body. However, given that billions of cells die every day, many pre-apoptotic cells in a genetically unstable state may lurk in the body which could be susceptible to transformation in response to circulating cancer cfChPs. Systemic cancer treatments may also generate many cells that are karyotypically unstable and prone to transformation. Nonetheless, the relevance of our findings to real world metastasis would require future clinical validation.

In conclusion, our study is the first to demonstrate that somatic cells can be transformed into a pluripotent state by naturally occurring agents in the form of circulating cfChPs. Clone D5 cells showed upregulation of the stem cell markers that have until now been achieved by artificial means by the introduction of stem cell transcription factors using viral vectors^33^. Our results also suggest that cfChPs are naturally occurring oncogenic agents with implications in cancer metastasis. We believe that clone D5 may be a valuable resource of further research into naturally occurring mechanisms of oncogenesis and pluripotency.

## Materials and Methods

### Ethics approval (human subjects)

This study was approved by Institutional Ethics Committee (IEC) of Advanced Centre for Treatment, Research and Education in Cancer (ACTREC), Tata Memorial Centre (TMC) for collection of blood from cancer patients for isolation of cfChPs (Approval no. 900011). All participants signed a written informed consent form approved by the IEC.

### Development of clone D5

The single cell clone D5 was developed in the year 2011 from NIH3T3 cells treated with cfChPs isolated from pooled sera of cancer patients.

#### Isolation of cfChPs from serum

Blood was collected from cancer patients (10mL) in plain vacutainers and was allowed to clot. Serum was separated, and serum from 5 patients was pooled for isolation of cfChPs. Demographic details of the 5 cancer patients from whom blood was collected is given in Supplementary Table S5. cfChPs were isolated according to a protocol described by us in detail earlier^21^. The steps involved in the isolation protocol are briefly summarised herewith: 1) ultracentrifugation of 1 ml aliquots of the pooled serum at 700,000g for 16 h at 4°C; 2) treatment of the pellet obtained with lysis buffer followed by repeat ultracentrifugation as above; 3) suspending the pellet obtained in 1 ml PBS and passing the suspension through an affinity column (ThermoFisher Scientific, USA) containing biotinylated anti-histone H4 antibody (125 µg) bound to 2 mL of Pierce® Streptavidin Plus Ultralink® Resin (ThermoFisher Scientific, USA); 4) elution of the column with 1 mL 0.25 M NaCl followed by ultra-centrifugation of the elute as described above; 5) suspending the pellet containing cfChPs in 1 mL PBS. The concentrations of cfChPs in the isolates are expressed in terms of their DNA content, as estimated using the PicoGreen dsDNA quantitation assay (ThermoFisher Scientific, USA). A representative electron microscopy image of the isolated cfChPs is given in Supplementary Figure S7.

#### Treatment of NIH3T3 cells with cfChPs

NIH3T3 mouse fibroblast cells (CRL-1658) were obtained from the American Type Culture Collection (ATCC) and maintained in Dulbecco’s Modified Eagle’s Medium (DMEM) supplemented with 10% Bovine Calf Serum (BCS) in 35mm culture dishes at 37°C in humidified atmosphere containing 5% CO_2_. One hundred thousand NIH3T3 cells were treated with cfChPs (10ng) and the cells were passaged every fourth day. After the sixth passage, single cell clones were developed from the cfChPs treated cells by serial dilution method, and one of the clones (D5) was used in experiments described herein.

### Spheroid formation assay

Cells of D5 clone and NIH3T3 control cells were maintained in DMEM as described above. D5 and NIH3T3 control cells were suspended in 100 μL of MammoCult basal medium with proliferation supplement (STEMCELL Technologies, Inc., Vancouver, BC, Canada). One thousand cells were seeded in each well of ULA 96-well round-bottomed plates (Corning B.V. Life Sciences, Amsterdam, The Netherlands). The plates were incubated at 37°C in humidified atmosphere of 5% CO2. Images of spheroids were captured on bright field microscope at 20X magnification daily until day 7. The size of the spheroid was quantified using ImageJ software, and those with a diameter exceeding 60mm were visually counted.

### Flow cytometry

Flow cytometry was performed on NIH3T3 and D5 cells as well as on cells of the spheroids. For the former, cells were harvested by gently scraping the cells using a cell scraper and placed in FACS tubes. For the latter, spheroids from multiple wells were harvested and trypsinized into a single cell suspension. Cells were then stained with the stem cell antibodies CD34 PE, CD133 FITC, and CD44 APC for 20 minutes in dark followed by washing with 1X PBS. Cells were acquired on Attune NxT flow cytometer (Thermo Fisher Scientific) and data were analysed using FlowJo™ Software version 10.6 (BD, Ashland, OR, USA). Cells were gated based on forward and side scatter characteristics and a minimum of 20,000 cells were acquired. The median fluorescence intensity (MFI) was then compared between control NIH3T3 cells and those of D5 clone.

### qRT-PCR

The stem cell transcription factors were assessed on NIH3T3 and D5 cells as well as on cells of the spheroids using quantitative reverse transcription polymerase chain reaction (qRT-PCR). Total RNA was extracted using RNeasy Mini Kit (Qiagen, Hilden, Germany) and was converted to complementary DNA (cDNA) using the High Capacity cDNA Reverse Transcription Kit with RNase Inhibitor (Applied Biosystem, CA, USA). cDNA (∼12.5ng) was used for qRT-PCR to estimate mRNA expression of the stem cell transcription factors OCT4, NANOG, and SOX2, utilizing specifically designed primer sets. Real-time PCR amplification was carried out using SYBR Select Master Mix (Applied Biosystems, CA, USA), with samples assayed on a QuantStudio™ 5 Real-Time PCR System (Thermo Fisher Scientific, USA) utilizing a 384-well block in duplicates. Data analysis employed the comparative threshold cycle (CT) method, facilitating the calculation of fold change in mRNA expression as 2^(-ΔΔCT).

### Transcriptome analysis

#### RNA-seq quality control, mapping and quantification

The quality of raw FASTQ files was assessed using FastQC tool (v0.12.1)^52^, to ensure high-quality data and visualize using multiQC (v1.15)^53^. Salmon (v1.5.1)^54^ was used to map reads to the mouse reference transcriptome in a quasi-mapping mode, enabling accurate transcript abundance quantification with GC bias correction enabled. Transcript abundances were then transformed into gene-level counts via tximport (v1.18.0) package in R^55^ and DESeq2 (v1.30.1)^56^ was then applied to identify differentially expressed genes (DEGs). Subsequently, DEG results were visualised through a volcano plot using EnhancedVolcano^57^. Finally, heatmap of the DEG data was generated using pheatmap function of pheatmap package in R.

#### Enrichment Analysis

The software GSEA; v4.3.3^58^ was used to perform pathway enrichment analysis on DESeq2-normalized counts using msigdb.v2023.2.Mm.symbols.gmt gene set. One thousand phenotype permutations were conducted and ‘diff_of_classes’ ranking metric was used to generate the ranked list of genes. Enrichment was assessed based on normalised enrichment score (NES), where positive scores indicate upregulation and negative scores indicate down-regulation. A nominal (nom) p-value of less than or equal to 0.05 was considered statistically significant.

GeneSetCluster in R (available at https://github.com/TranslationalBioinformaticsUnit) was employed to cluster pathways based on shared genes^59^. The distance between gene sets was computed using the Jaccard index. Subsequently, the kmeans method was applied to cluster the gene sets into groups based on calculated distance. To ascertain the optimal number of clusters, OptimalGeneSets was utilized using gap method.

### Animal Ethics approval

The experimental protocol of this study was approved by the Institutional Animal Ethics Committee (IAEC) of the Advanced Centre for Treatment, Research and Education in Cancer (ACTREC), Tata Memorial Centre (TMC). The animal study (SCID mouse inoculation described below) was conducted twice, once in 2018 (IAEC approval no. 12/2018) and in 2024 (IAEC approval no. 19/2024). The experiments were carried out in compliance with the IAEC animal safety and ARRIVE guidelines.

ACTREC-IAEC maintains respectful treatment, care and use of animals in scientific research. It aims to use animals in research for advancement of knowledge following the ethical and scientific necessities. All scientists and technicians involved in this study have undergone training in the ethical handling and management of animals under the supervision of FELASA certified attending veterinarian.

All welfare considerations to minimize animal suffering and distress were undertaken. Mice were observed once every two days for the humane endpoints, viz. tumor size and activity and weight loss of more than 15 %. None of the animals were found to have reached the above humane endpoints at the time of sacrifice.

### Inoculation of D5 cells into SCID mice

Inbred female NOD SCID (NOD.Cg-Prkdcscid/J) mice were obtained from the institutional animal facility and maintained according to IAEC standards. They were housed under maximum-barrier facilities in sterilized cages with filter tops and sterile bedding material. The mice were fed γ-irradiated sterilized food and water ad libitum and kept under 12 h light/dark cycle with free access to water and food. A HVAC system provided controlled room temperature, humidity and air pressure. The NOD SCID mouse inoculation study was conducted twice: in 2018 and in 2024. In both cases, one million D5 clone cells and an equal number of NIH3T3 control cells were inoculated in mice. In the first experiment, 5 mice each were inoculated in the control and experimental groups, while in the second experiment, 9 control mice and 8 experimental mice were inoculated.

### Immuno-histochemistry

Immuno-histochemistry (IHC) to detect germ cell markers was done on the D5 induced tumours in both the experiments on formalin fixed paraffin embedded (FFPE) sections as per standard procedure. Tissues were stained with primary antibodies against the germline cell markers, viz. ectoderm: [glial fibrillary acidic protein (GFAP) (Dako, Denmark)]; mesoderm: [CD45 (Abcam, Cambridge, UK)] and endoderm: [alpha fetoprotein (AFP) (GeneTex, Irvine, California)]. The IHC Select® HRP/DAB 150 kit (Merck KGaA, Darmstadt, Germany) was used as the secondary antibody to detect colour development.

### FISH analysis

FISH analysis were done on metaphase spreads from D5 cells and on unstained FFPE sections of tumours that had developed in SCID mice. FISH was performed using human specific whole-genomic probe (Applied Spectral Imaging, Israel) as described by us earlier^21^. Images for both metaphase spreads and tumor sections were acquired under Applied spectral bio-imaging system (Applied Spectral Imaging, Israel) and examined for the presence of human DNA signals using FISH-View software 8.0 (Applied Spectral Imaging, Israel).

## Acknowledgements

We thank Mr. Rohan Chaubal for his inputs while writing the manuscript and Dr. Kavita Pal for supervising one of the animal inoculation experiments. Transcriptome sequencing was outsourced to Medgenome Pvt. Ltd. India.

## Author Contributions

I.M. conceptualized the study; S.S., N.L.D., S.M., R.K., R.L., N.A., G.V.R., N.K.K., V.J. and S.Shinde. performed the experiments; S.S., N.L.D. and P.C. performed data analysis; I.M. and P.C. supervised the study; I.M. procured funding; I.M., S.S., N.L.D. and P.C. wrote the manuscript; I.M. approved the final draft.

