## Supplementary material for "Induction of pluripotent oncogenic stem cells from mouse fibroblasts": Legends to Figures and Supplementary Figures

**Figure 1: Induction of pluripotent stem cells from NIH3T3 cells.** **(a)** Upregulation of mRNA expression of stem cell-related transcription factors NANOG, OCT4, and SOX2 in D5 clone cells and NIH3T3 cells as detected by qRT-PCR and analysed using a comparative C_T_ method. **(b)** Flow cytometry plots and adjacent box plots represent the relative upregulation of CD34 and CD133 expression in the D5 clone cells (red) compared to the NIH3T3 control cells (blue). The degree of upregulation of CD44 observed in D5 clone was not statistically significant from NIH3T3 cells. **(c)** Representative images showing growth kinetics and morphological changes of spheroids formed by D5 clone and NIH3T3 control cells over 7 days in culture. **(d)** Box plots representing the increased number and size of spheroids formed by D5 clone cells compared to NIH3T3 cells on day 7. **(e)** Upregulation of mRNA expression of stem cell-related transcription factors NANOG and OCT4 in spheroid cells of D5 clone compared to those of NIH3T3 cells as detected by qRT-PCR and analysed using a comparative C_T_ method. **(f)** Representative flow cytometry plots and adjacent respective box plots to represent the relative upregulation of CD34, CD133 and CD44 expression in spheroid cells of D5 clone (red) compared to NIH3T3 control cells (blue). The box plots represent the median with minimum and maximum values. Statistical analyses were performed using two-tailed unpaired student’s *t*-tests in GraphPad Prism 8. **_*_** p<0.05, **_**_** p<0.01, **_****_** p<0.0001, ns non-significant.

**Figure 2: RNA-seq data quality assessment and differential expression analysis.** **(a)** Per sequence quality scores **(b)** MDS plot shows patterns of clusters of control and D5 samples **(c)** Enhanced volcano plot of DEGs. Red nodes represented up-regulated genes with log 2 FC > 1.5 and adjusted p < 0.05; blue nodes represented down-regulated genes with log 2 FC < -1.5 and adjusted p < 0.05.

**Figure 3: Significantly enriched pathways and gene clusters in D5 samples.** **(a)** The enriched pathways demonstrating a significant p-value < 0.05 and NES > 1.5 and highlighted those acquiring cancer stem cell-like characteristics **(b)** GeneSetCluster plot showing relative distance scores of significantly enriched pathways.

**Figure 4: Subcutaneous inoculation of cells of D5 clone generate pluripotent tumours. (a)** SCID mouse harboring a tumour; **(b)** H&E section of the tumour show fibro-sarcoma; **(c)** FISH analysis on FFPE section of a tumour reveal presence of human DNA; **(d)** Representative IHC images showing expression of germline markers in FFPE sections of a tumour. GFAP = Glial fibrillary acidic protein, CD45 = leukocyte common antigen, AFP = alpha-fetoprotein.

**Legends to Supplementary Figures**

**Supplementary Figure S1: Morphological changes of D5 cells overtime and presence of human DNA in them. (a)** Representative images of NIH3T3 cells and those of clone D5 in 2014 and 2024. **(b)** FISH analysis on metaphase spreads prepared from D5 cells in 2024 show human DNA to be associated with the chromosomes.

**Supplementary Figure S2:** Flow cytometry plots representing the gating strategy based on forward and side scatter parameter.

**Supplementary Figure S3: Quality control analysis of RNA-Seq data. (a)** Mean quality scores of RNA-seq data **(b)** Cook’s distance boxplot calculated for each sample **(c)** Heatmap showing significant DEGs.

**Supplementary Figure S4: Upregulated pathways in D5 samples. (a)** Hallmark P53 pathway **(b)** Biocarta CTCF pathway **(c)** Biocarta ATM Pathway **(d)** WP Apoptosis **(e)** WP ERBB signaling pathway and **(f)** Reactome Oncogene Induced Senescence. These plots feature vertical black lines representing genes within gene sets on x-axis and ES on y-axis. A green line connects ES points to corresponding genes indicating the degree of over-representation. Colored band indicate positive or negative correlation. NES and nominal p-value were provided for each gene set to assess the significance of enrichment.

**Supplementary Figure S5: Downregulated pathways in D5 samples. (a)** Reactome Respiratory electron transport **(b)** Reactome the citric acid TCA cycle and respiratory electron transport, **(c)** Reactome Chemokine receptors and chemokines and **(d)** Reactome class A I Rhodopsin like receptors. These plots feature vertical black lines representing genes within gene sets on x-axis and ES on y-axis. A green line connects ES points to corresponding genes indicating the degree of over-representation. Colored band indicate positive or negative correlation. NES and nominal p-value were provided for each gene set to assess the significance of enrichment.

**Supplementary Figure S6: Geneset clusters in D5 samples.** Heatmap plot showing relative distance scores of significantly enriched pathways. The list of genes and pathways is also shown in the plot. Five distinct clusters of pathways are highlighted.

**Supplementary Figure S7: EM image of cfChPs isolated from pooled sera of cancer patients.** The image shows a bead-on-a-string appearance typical of chromatin. Reproduced with permission from ref 21.
