## Supplementary Table S5 for "Induction of pluripotent oncogenic stem cells from mouse fibroblasts"

**Details of cancer patients**

| **Sr. No.** | **Age** | **Sex** | **Diagnosis** | **Stage** |
| --- | --- | --- | --- | --- |
| 1. | 31 Years | Female | Ca Breast | IV |
| 2. | 68 Years | Male | Ca Lung | IV |
| 3. | 47 Years | Male | Ca Lung | IV |
| 4. | 44 Years | Female | Ca Breast | III |
| 5. | 58 Years | Male | Ca Esophagus | III |
